# Evaluating the Survival and Removal of *Escherichia coli* from Surfaces Made with Traditional and Sustainable Cement-Based Materials in Field-Relevant Conditions

**DOI:** 10.1101/2024.11.07.622504

**Authors:** Claire E. Anderson, Jason Hernandez, Suhi Hanif, Lauren Owens, Yoshika Crider, Sarah L. Billington, Michael Lepech, Alexandria B. Boehm, Jade Benjamin-Chung

## Abstract

Soil household floors are common in low- and middle-income countries (LMICs) and can serve as reservoirs of enteric pathogens. Cement-based floors may interrupt pathogen transmission, but little is known about pathogen survival or removal from cement-based surfaces. This study investigated the survival of *Escherichia coli* (*E. coli*), an indicator of fecal contamination, on cement-based surfaces and evaluated its reduction through common household activities (mopping, sweeping, and walking). We compared *E. coli* fate on three mixes: 1) Ordinary Portland Cement (OPC) concrete (used in the United States), 2) OPC mortar (used in Bangladesh), and 3) OPC mortar with fly ash (a sustainable alternative to the Bangladesh mix). Additionally, we compared outcomes on cement-based surfaces with and without soil and at two temperatures representing the dry and wet seasons in Bangladesh. After 4 hours on the cement-based surfaces, *E. coli* decayed more than 1.1 log_10_(*C*/*C*_*o*_) under all conditions tested, which is significantly faster than in bulk soils. The higher temperature increased the decay rate constant (p = 5.56*10^−8^) while soil presence decreased it (p = 2.80*10^−6^). Sweeping and mopping resulted in high levels of removal for all mixes, with a mean removal of 71% and 78%, respectively, versus 22% for walking. The concrete and mortar mix designs did not impact *E. coli* survival or removal (p > 0.20). Cement-based floors made with a fly ash mix performed similarly to traditional cement-based floors, supporting its potential use as a more sustainable intervention to reduce fecal contamination in rural LMIC household settings.

**Importance:** Cement-based surfaces may serve as a health intervention to reduce the fecal-oral transmission of pathogens in household settings, but there is a critical lack of evidence about the fate of indicator organisms on these surfaces, especially in field-relevant conditions. This study provides some of the first insights into *E. coli* survival on cement-based surfaces and the effectiveness of daily activities for removing *E. coli*. Additionally, this study explores the fate of *E. coli* on cement-based surfaces made with fly ash (which contributes fewer CO_2_ emissions) versus traditional cement mixes. We found that *E. coli* had similar survival and removal across all mix designs, demonstrating that fly ash mixes are feasible for use in household settings (e.g., in floors). The findings enhance understanding of fecal-oral transmission pathways and support the use of fly ash mixes in cement-based flooring in future epidemiologic studies assessing effects on enteric disease burdens.

## Introduction

Enteric pathogens cause gastrointestinal issues including stomach cramps, bloating, nausea, and diarrhea, and have been linked to growth stunting, malnutrition, impaired child development, and mortality (1, 2). Worldwide, diarrhea was the third leading cause of death among children 5 years and younger in 2024 and was responsible for 59.3 million disability-adjusted life years in 2021 (3, 4). Enteric infections are especially prevalent in low- and middle-income countries (LMICs) in South Asia, Southeast Asia, and Sub-Saharan Africa, where transmission commonly occurs via fecal-contaminated hands, food, water, household surfaces, and/or soil (1, 5). Enteric pathogens that cause diarrhea can often survive and reproduce in soil, causing it to act as a reservoir of disease (5–8). Children can ingest soil contaminated with human or animal feces while crawling and playing, leading to exposure to pathogens and potential infection (5, 7, 9–12). Soil household floors are extremely common in rural LMIC settings and represent an environmental transmission pathway within households (13). Although medical interventions may be able to successfully treat enteric infections, continued exposure to contaminated environments limits the ability for infection control.

Replacing soil floors with finished floors, such as concrete or mortar, is a promising intervention aimed at eliminating soil reservoirs for pathogens and interrupting environmentally mediated transmission pathways. Finished flooring may facilitate pathogen removal by being easier to clean than soil flooring, as pathogens cannot be fully removed from soil floors. Previous research studies in Bangladesh and Kenya have shown that houses with soil floors have a higher prevalence of soil-transmitted helminths and *Giardia* infections than households with finished floors (14, 15). In Mexico, a large-scale government program replacing dirt floors with concrete flooring resulted in observed decreases in parasitic infestations, diarrhea, a lower prevalence of anemia, and an improvement in the cognitive development of treated children based on various surveys and biological samples (16). Additional studies (17–19) have demonstrated that living in a household with finished floors is associated with lower disease risk and higher well-being. An ongoing randomized trial, Cement flooRs AnD chiLd hEalth (CRADLE, NCT05372068), in rural Bangladesh is evaluating whether replacing household soil floors with cement-based floors reduces soil-transmitted helminth infections and diarrhea in children under 2 years (20). While observational studies have demonstrated that living in a household with finished floors is associated with lower disease risk and higher well-being, the underlying mechanisms for these improvements remain unclear.

A critical, but often overlooked consideration if cement-based floors were to be scaled up in LMICs as a health intervention is the environmental impact, as cement production contributes an estimated 5 to 10% of total anthropogenic CO_2_ emissions (21, 22). The Lancet Commission on planetary health named the development of interventions that improve health while minimizing greenhouse gas emissions as a research priority (23). To mitigate this impact, the CRADLE study is unique in that it examines the use of a supplementary cementitious material (SCM), fly ash, to create an alternative cement mix that may be produced with fewer CO_2_ emissions. Fly ash is a by-product of coal combustion in coal-fueled power plants. It is commonly used in the construction industry (21) and is readily available in markets in LMICs. Using fly ash as a replacement for cement can potentially reduce the impact of CO_2_ emissions by as much as 40% depending on the replacement level (24–31). In addition, the structural impact of incorporating fly ash in concrete mixes has been rigorously tested, demonstrating improved strength and durability of the concrete compared to traditional concrete and mortar mixes (26–31). However, in principle, it is possible that differences in each cement mix’s chemical properties, surface roughness, porosity, pH, moisture retention, or other factors could impact the survival and removal of pathogens on each mix’s surface. For example, increased surface roughness could provide protection of *Escherichia coli* (*E. coli*) from removal activities or environmental stressors. If the fly ash cement mix increased the duration of pathogen survival and decreased its removal during cleaning or other common household activities, it could support higher levels of pathogen transmission, which may outweigh any environmental benefits with respect to CO_2_ emissions.

Indeed, the survival of indicators of contamination and enteric pathogens may be impacted by numerous factors that affect the desiccation of the pathogen, including temperature, humidity, and organic matter presence (32–40). However, to our knowledge, no prior studies have investigated the removal efficiency of *E. coli* from cement-based surfaces nor the removal efficiency of common household activities. Studies investigating the removal of *E. coli* from surfaces focused on chemical removal methods, again from non-porous surfaces (41–44). Studies investigating *E. coli* persistence on surfaces have primarily focused on food manufacturing environments and non-porous surfaces like glass, plastic, or metal (37, 38, 45–49). Similarly, studies investigating soil are limited to agricultural soils with manure application (39, 50–52), or large volumes of soil rather than dust or dirt on the surface of floors (53–56). One prior study investigated the survival of *E. coli* on concrete, however, the temperature was not constant in the study and ranged from 20 to 28 °C (57). Generally, this study found that *E. coli* was less persistent on concrete versus sand or gravel (57).

We hypothesize that the alternative chemical properties and differing surface roughness between cement mixes are not substantial enough to alter the survival or removal of enteric pathogens on cement surfaces using different mix designs during common household activities. We investigated this hypothesis using the fate of *E. coli* on one concrete and two mortar mix designs to assess whether the health impact of a cement-based floor intervention may be influenced by the choice of mix designs. *E. coli* serves as an indicator of fecal contamination (58, 59) and is a biosafety level 1 pathogen surrogate to enteric pathogens that is practical to work with in laboratory settings. In this study, we used field-relevant conditions to the CRADLE trial in Bangladesh to assess the survival of *E. coli* on three surfaces: an Ordinary Portland Cement (OPC) concrete mix typically used in the United States, an OPC mortar mix typically used in Bangladesh, and an OPC mortar mix with 25% class F Fly Ash as a sustainable alternative (referred to as OPC fly ash mortar in this study). We chose the inoculum used to spike the *E. coli* on the cement-based surfaces, the timeline of sampling, temperature, humidity, soil, and removal activities based on what is common to Bangladesh. Understanding the survival and removal of *E. coli* on cement-based surfaces addresses a critical knowledge gap in the literature and can provide insights into the mechanisms through which cement-based floors may reduce the transmission of fecal-oral pathogens in rural LMIC settings.

## Results

All negative controls from the cement-based tiles (127 mm squares with 12.7 mm depth to mimic household cement-based floors in the laboratory) were verified to have zero CFU, and positive controls had consistent *E. coli* concentrations (∼10^7^ CFU/mL). Approximately 3.6% of *E. coli* was recovered from the cement-based tile squares, with mean recoveries of 3.4% from the OPC concrete, 4.2% from OPC mortar tiles, and 2.8% from OPC fly ash mortar. The standard deviations of these mean recoveries overlapped across the different mix designs, indicating mix design did not affect recovery. With soil, recovery was approximately 3.5%, and without soil, recovery was approximately 3.7%, again indicating little effect on recovery. In terms of the roughness profile results, OPC concrete had the highest average roughness at 55.8 μm, followed by OPC mortar at 30.2 μm, and then OPC fly ash mortar at 25.2 μm (**Table 1**). When comparing the average roughness, no mix design was substantially smoother than another, as all average roughness values were within one standard deviation of one another.

**Table 1:** Surface roughness profile information for the three mix designs.

| Mix Design | Average Roughness ±<br>Standard Deviation<br>($\mu\text{m}$) | Root-Mean-Square<br>Roughness ±<br>Standard Deviation<br>($\mu\text{m}$) | Total Roughness Height<br>± Standard Deviation<br>($\mu\text{m}$) |
| --- | --- | --- | --- |
| OPC Mortar | 30.2 ± 19.8 | 36.6 ± 22.2 | 149.7 ± 34.3 |
| OPC Concrete | 55.8 ± 13.1 | 68.5 ± 14.3 | 247.9 ± 41.1 |
| OPC Fly Ash Mortar | 25.2 ± 11.0 | 29.3 ± 13.1 | 85.6 ± 27.1 |

In the survival experiments, all decay rate constants were significantly different from zero, indicating decay of the bacteria with time (p < 4.5*10^−5^ across the 12 regressions, **Table 2**). Decay rate constants varied from 0.01 min^-1^ to 0.06 min^-1^ and represent the slope of a log-linear regression model run for each condition across the three trials (12 regressions in total, **Table 2**). ANOVA results from comparing the decay rate constants of the 36 experiments (12 conditions x 3 trials) indicate that mix design had no association with the survival of *E. coli* on the cement-based tiles (p = 0.20). In contrast, the environmental condition and the presence of soil were significantly associated with the decay rate constant. Post-hoc tests indicated higher temperatures were associated with faster decay (p = 5.56*10^−8^, mean decay rate constant difference of 0.019 min^-1^) compared to lower temperatures, and the presence of soil was associated with slower decay (p = 2.80*10^−6^, mean decay rate constant difference of 0.016 min^-1^) compared to the absence of soil. The arithmetic mean of the untransformed data, when converted to log_10_, showed an approximate reduction across mix designs after 2 hours of 2.0 log_10_(*C*/*C*_*o*_) with soil in the wet season, 1.3 log_10_(*C*/*C*_*o*_) with soil in the dry season, 2.4 log_10_(*C*/*C*_*o*_) without soil in the wet season, and 0.90 log_10_(*C*/*C*_*o*_) without soil in the dry season. In general, the highest concentration reduction for a single condition after two hours was achieved with the OPC mortar mix in the wet season without soil (3.4 log_10_(*C*/*C*_*o*_)) (**Figure 1**). After 4 hours, the *E. coli* concentration at this condition for all mix designs was below our limit of detection. In contrast, the lowest concentration reduction after two hours occurred with the OPC fly ash mortar in the dry season with soil (0.77 log_10_(*C*/*C*_*o*_)). After 4 hours in this condition, the mean reduction across mix designs was 1.3 log_10_(*C*/*C*_*o*_), while it was 2.3 log_10_(*C*/*C*_*o*_) with soil in the wet season, and 1.9 log_10_(*C*/*C*_*o*_) without soil in the dry season. The complete survival experiment results showing the natural log reduction in concentration versus time are plotted in **Figure 1**.

**Table 2:**
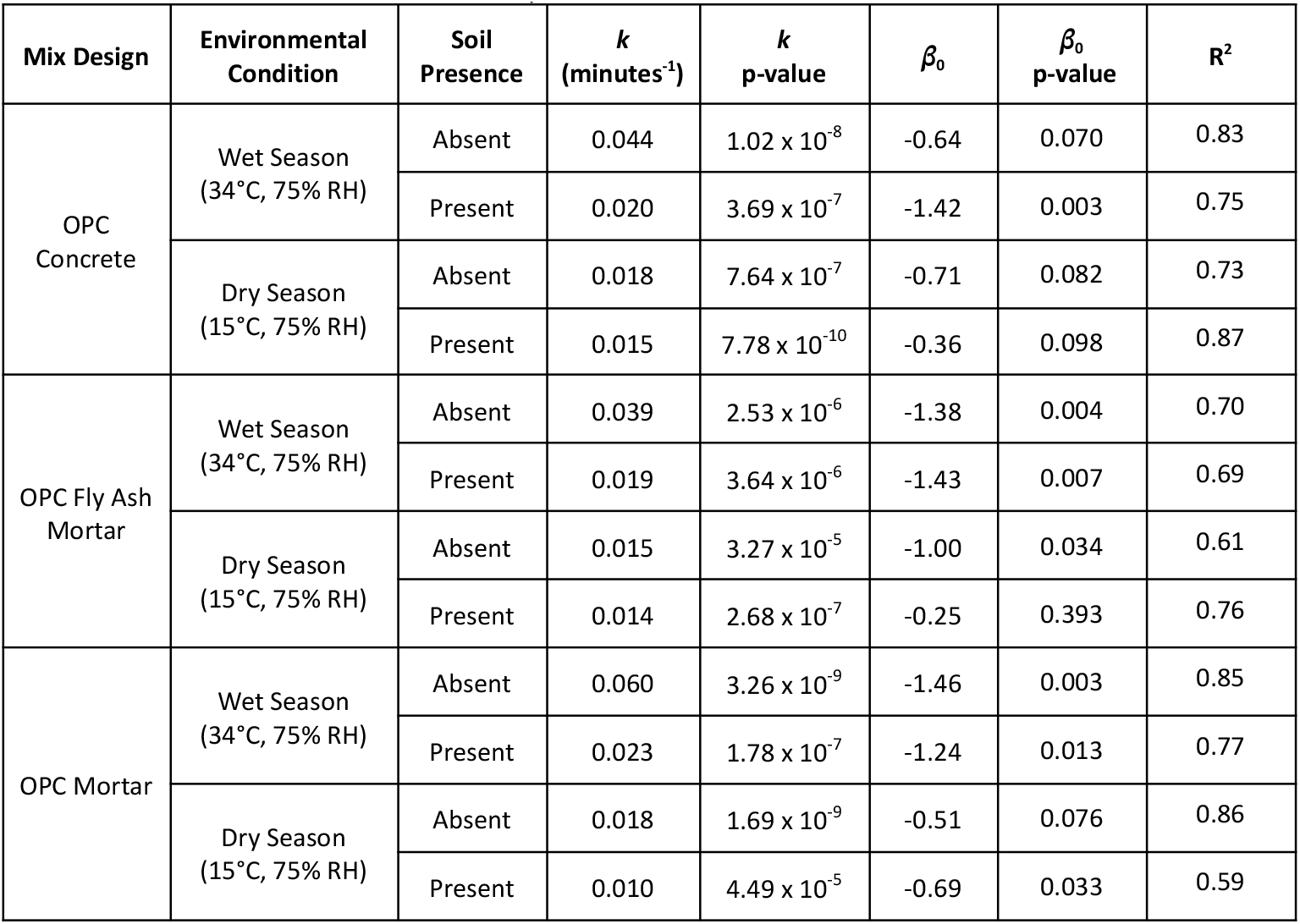
Decay rate constants (*k*) and y-intercept values (*β*_0_) from regression analyses for each experimental condition.

| Mix Design | Environmental Condition | Soil Presence | $k$<br>(minutes<sup>-1</sup>) | $k$<br>p-value | $\beta_0$ | $\beta_0$<br>p-value | R<sup>2</sup> |
| --- | --- | --- | --- | --- | --- | --- | --- |
| OPC Concrete | Wet Season<br>(34°C, 75% RH) | Absent | 0.044 | $1.02 \times 10^{-8}$ | -0.64 | 0.070 | 0.83 |
| | | Present | 0.020 | $3.69 \times 10^{-7}$ | -1.42 | 0.003 | 0.75 |
| | Dry Season<br>(15°C, 75% RH) | Absent | 0.018 | $7.64 \times 10^{-7}$ | -0.71 | 0.082 | 0.73 |
| | | Present | 0.015 | $7.78 \times 10^{-10}$ | -0.36 | 0.098 | 0.87 |
| OPC Fly Ash Mortar | Wet Season<br>(34°C, 75% RH) | Absent | 0.039 | $2.53 \times 10^{-6}$ | -1.38 | 0.004 | 0.70 |
| | | Present | 0.019 | $3.64 \times 10^{-6}$ | -1.43 | 0.007 | 0.69 |
| | Dry Season<br>(15°C, 75% RH) | Absent | 0.015 | $3.27 \times 10^{-5}$ | -1.00 | 0.034 | 0.61 |
| | | Present | 0.014 | $2.68 \times 10^{-7}$ | -0.25 | 0.393 | 0.76 |
| OPC Mortar | Wet Season<br>(34°C, 75% RH) | Absent | 0.060 | $3.26 \times 10^{-9}$ | -1.46 | 0.003 | 0.85 |
| | | Present | 0.023 | $1.78 \times 10^{-7}$ | -1.24 | 0.013 | 0.77 |
| | Dry Season<br>(15°C, 75% RH) | Absent | 0.018 | $1.69 \times 10^{-9}$ | -0.51 | 0.076 | 0.86 |
| | | Present | 0.010 | $4.49 \times 10^{-5}$ | -0.69 | 0.033 | 0.59 |

**Figure 1:**
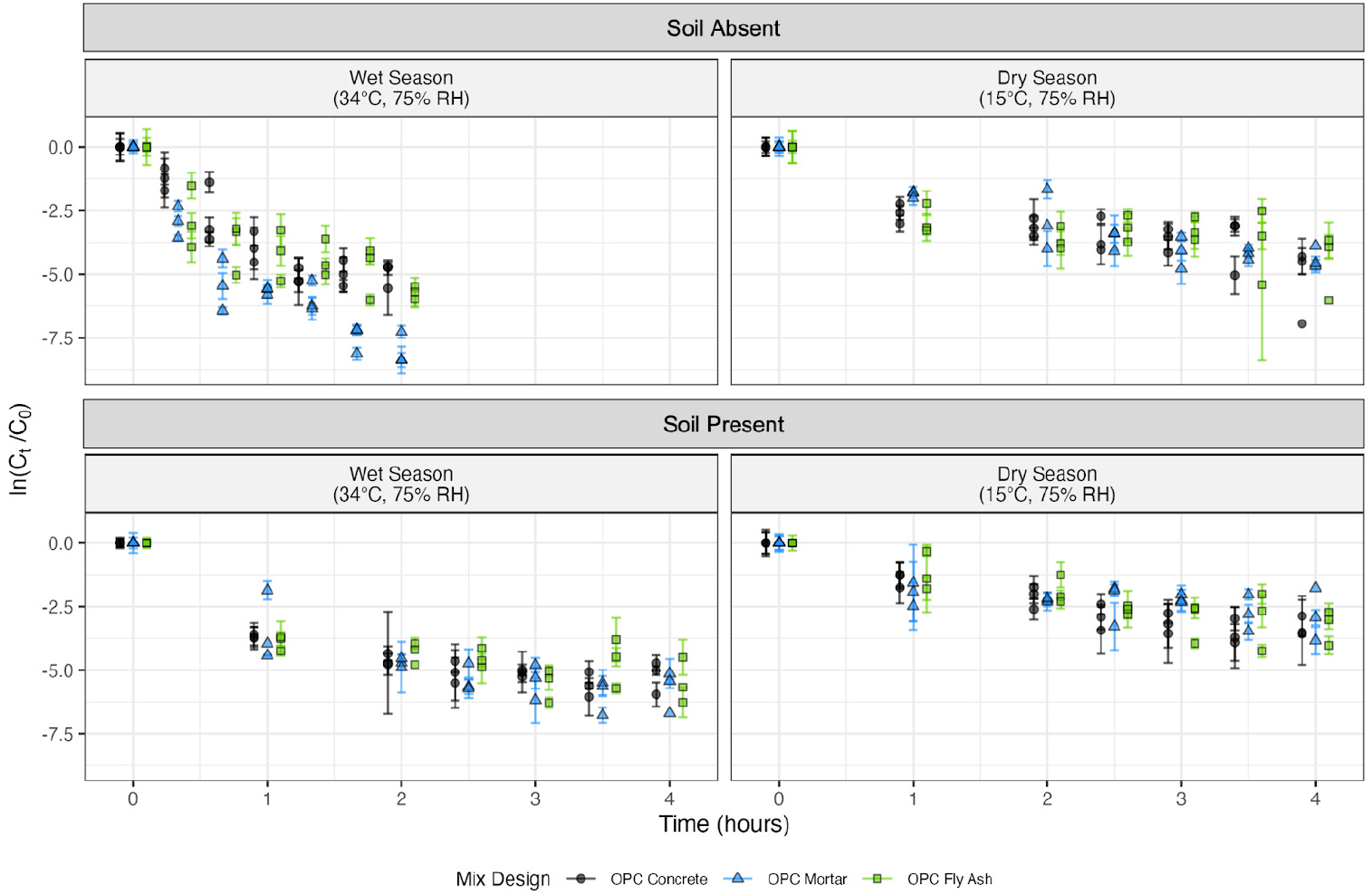
Natural log reduction of *E. coli* versus time (hours). The data are split into subplots based on the presence of soil and the environmental conditions for the incubation of the cement-based tile. The average reductions in concentration are plotted for each trial and the three mix designs,with error bars representing the 95% confidence interval for the reduction based on the CFU counts within each trial. Data points are jittered to better visualize the differences between the mix designs.

The effectiveness of *E. coli* removal varied significantly depending on the removal activity (p = 3.9*10^−10^) and the presence of soil (p = 3.4*10^−3^), while the mix design was not a significant factor controlling removal efficiency (p = 0.34). The average percent of *E. coli* removed by mopping was 88% for OPC concrete with soil, 74% for OPC mortar with soil, and 80% for OPC fly ash mortar with soil (**Figure 2**). Without soil, the mopping removal was 67%, 82%, and 75% for OPC concrete, OPC mortar, and OPC fly ash mortar, respectively. For sweeping, the average *E. coli* removal without soil was 50%, 46%, and 65% for OPC concrete, OPC mortar, and OPC fly ash mortar, respectively. Average sweeping removal with soil was higher than without, at 82% for OPC concrete, 93% for OPC mortar, and 89% for OPC fly ash mortar. Finally, the average *E. coli* removed from walking without soil was 51% for OPC concrete, 25% for OPC mortar, and 37% for OPC fly ash mortar. Walking removal was lower (9%, 4%, and 9% for OPC concrete, OPC mortar, and OPC fly ash mortar, respectively) with soil. Post-hoc tests showed that the *E. coli* removal from mopping and sweeping was significantly more than the *E. coli* removal from walking (p = 1.7*10^−9^ and p = 6.0*10^−8^, respectively and a mean removal difference of 55% and 49%, respectively), while the removal values from sweeping and mopping were not significantly different from one another (p = 0.56). The two-way interaction between removal activity and the presence of soil (p = 1.2*10^−5^) was found to be significant, with a post hoc test revealing that *E. coli* removal was higher with soil for sweeping (p = 7.5*10^−6^, mean removal difference of 34%), but did not significantly impact mopping or walking (p = 0.76 and p = 0.48, respectively).

**Figure 2:**
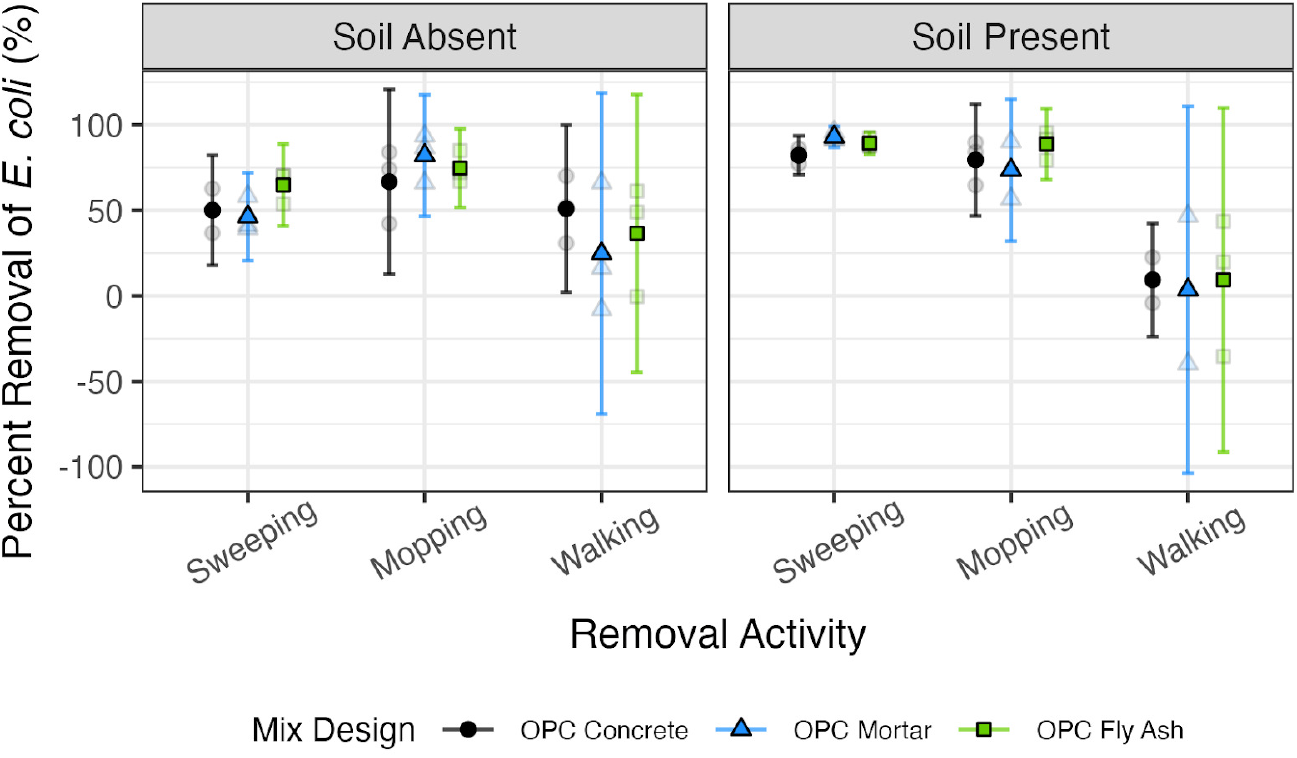
Percent removal of *E. coli* from cement given three removal activities. The data are split into subplots based on the presence of soil. The average percent removal for each of the three trials (opaque points) and the percent removal from each trial (transparent points) are plotted, in addition to error bars representing the 95% confidence interval for the trials. Mix designs are ordered in decreasing average roughness measurements.

## Discussion

We found that OPC concrete (typically used in the United States), OPC mortar (typically used in rural Bangladeshi homes), and OPC fly ash mortar (which represents a mix with lower embodied carbon), had similar survival and removal efficiencies for *E. coli*. Similarities in the fate of *E. coli* on the mix designs are important to validate because although the sustainable alternative of OPC fly ash mortar has been rigorously tested for similar mechanical and durability properties to traditional OPC mortar (25, 30), there has been little investigation into the microbial implications of this lower carbon material. The fate of *E. coli* may have been similar between the mixes due to negligible differences in the surface roughness between mixes, as surface roughness only differed by 20.4 μm on average. The observed similarities in *E. coli* fate between traditional cement-based mix designs and the alternative that incorporated fly ash indicate that enteric pathogen transmission will not be influenced by these mix designs, supporting the use of more sustainable cement-based materials using fly ash in future epidemiological studies to assess health impacts of cement-based household floors.

*E. coli* is an indicator of the persistence of fecal contamination and the possible survival of other pathogens. Studies investigating *E. coli* survival on non-porous, dry surfaces at room temperature reported decay rate constants generally less than 0.02 min^-1^, which are lower than those found in this study (38, 45–47, 60). In one study that analyzed decay on concrete sidewalks, *E. coli* decay was very similar to the measured decay in this study under similar conditions (57). Specifically, they observed an approximately 2 log_10_(*C*/*C*_*o*_) decay at a temperature ranging from 20 -28°C after two hours in darkness, versus our measured decay of 2.4 log_10_(*C*/*C*_*o*_) at 34°C after two hours. At approximately room temperature in the United States (∼21°C), decay rate constants for *E. coli* in sterile soil, agricultural soils, and soils with manure were less than 0.0002 min^-1^ (53–56, 61) versus the decay rates found in this study, which were >0.010 min^-1^. Given these results, cement-based flooring likely reduces the risk of human interaction with possible pathogens versus soil flooring, but several factors should be considered when applying these results to household flooring scenarios in LMICs. If we assume there are approximately 10^9^ CFU *E. coli*, representing 1 g of fecal contamination (62) on the cement-based surface, and we are able to recover 100% of the *E. coli* from the surface, the lowest decay rate constant measured in this study indicates that the *E. coli* would be undetectable (≤ 1 CFU) after about 35 hours. In contrast, with the highest decay rate constant from this study, *E. coli* would be undetectable after approximately 6 hours. In soil under these same conditions, the E. coli would take over 2 months to become undetectable. Household concrete flooring is likely to be disturbed within 6-35 hours, when indicator species are still detectable. Walking can transfer *E. coli* from the cement-based surface to shoes or human skin, enabling further spread within the household, to other households, or to community spaces (63). Furthermore, there are high levels of child contact with flooring which may take place within minutes of contamination, so although fecal indicator species decay faster on cement-based flooring compared to soil, finished flooring may not reduce risk completely (8, 64). Our results of *E. coli* survival and removal can be used together to estimate exposure to fecal contamination and possible pathogens on different household surfaces in quantitative microbial risk assessments (QMRAs) in future studies (65–67).

Notably in previous studies, *E. coli* survival was extremely variable, as it was detectable after hours, days, weeks, or sometimes months on non-porous surfaces at 4-12°C (32, 34, 37, 38, 48, 49). Large differences in *E. coli* survival times may be due to differences in *E. coli* strains. The decay of lab strains of *E. coli* may be different from the decay of environmental strains or pathogens, but previous studies have found mixed results of the impact of *E. coli* strain on its survival in soil (39, 52, 56, 58, 62). Differences in *E. coli* decay over time may also be due in part to biphasic decay. In biphasic decay, there is an initial greater decay rate in the first 24 hours as susceptible cells die off and a slower decay rate in the next several days as more robust cells persist (33, 37, 51, 68). This study intended to mimic CRADLE field conditions, where the flooring was unlikely to remain unperturbed for a long period of time. Thus, the survival experiments were limited to less than one day of measurements. By sampling for a short amount of time, we may have captured only the more rapid decay portion of a biphasic decay, if one exists.

*E. coli* decay rate constants are greater in the wet season conditions used in our study and are more pronounced when soil is absent, suggesting that temperature and protection from soil are important factors in decay. This finding agrees with previous studies which have investigated the decay of pathogens and fecal indicators and have shown that increasing temperature is positively correlated with increasing decay rate constants (32–40). Previous studies investigating the impact of soil (or other organic matter) presence on decay rate constants have generally found that soil increases the persistence of indicator species and pathogens on surfaces. Some studies have found that organic matter can have a barrier-like effect on organisms, protecting them from environmental stressors (33, 37). Additionally, sediment particle size can impact bacterial survival, with decay rate constants being less sensitive to temperature in finer soil (35). Soil composition, including clay content, had mixed effects on *E. coli* survival in previous studies (40, 53–56). In the field, we expect low levels of soils on the cement-based floors. In CRADLE pilot experiments, there was an average of 0.2 g/m^2^ (with a standard deviation of 0.3 g/m^2^) soil present on cement flooring (69). In this study, we used a larger mass of soil in a smaller area (equivalent to 25 g/m^2^ of soil) which may be representative of a spot of mud or soil tracked into the household rather than soil spread evenly across the cement-based surface. Our estimates of decay may be a lower bound for true household cement-based floor conditions, as the protective properties of soil may decrease with less soil. It will be important to note the density of dirt or soil on finished flooring in future field studies to assess its impact on bacterial decay.

Mopping and sweeping may not be as effective as chemical disinfectants, but our removal experiments indicate that these activities can effectively remove fecal contamination and possible pathogens from cement-based flooring surfaces. Results in this study ranged from 46 -93% removal efficiency after one activity and were statistically similar between the mix designs. Furthermore, while soil presence slowed *E. coli* degradation, it improved removal efficiency for sweeping. Improved removal efficiency could be because of adhesion of the *E. coli* to the soil, which then was removed in bulk through sweeping. Repeated cleaning throughout the day or the use of multiple removal activities (such as sweeping and then mopping) could be an effective preventive measure to reduce possible pathogens in the household (70). While chemical disinfectants such as hypochlorite solutions, ethanol, or peracetic acid can result in up to seven orders of magnitude reduction of *E. coli* within minutes (32, 41–44, 70–72), they are not field-relevant to CRADLE or similar studies as they are not commonly used in low-income rural LMIC households. Chemical disinfectants may not be used because they are expensive, unavailable in local markets, or may require a behavior change from individuals who clean the households (73–75).

This study contains limitations that can serve as potential topics for future work. First, this study focused on a single organism (*E. coli* K12), which is a fecal indicator bacteria. Fecal indicators, while useful, are limited in determining the potential presence of pathogens. Future work could focus on specific human pathogens (such as bacteria, viruses, or soil-transmitted helminths) for a setting of interest. Second, the study used a laboratory-cultured bacterial strain grown in high nutrient media in a stationary phase; environmental *E. coli*, which may be genetically diverse, may be in diverse growth states including senescence, causing its fate to diverge from that observed in this study. However, using lab strains is a necessary first step in experiments like this aiming to fill an important knowledge gap. Third, the cotton swabs, straw brooms, and rubber tiles that were used in the removal activities were used once to remove *E. coli* from the tile and then discarded. In reality, removal tools such as mops or brooms would be used multiple times. Previous studies have looked at sequential sampling methods (e.g. repeated finger touches on a surface) for microbial transfer (76, 77), and a similar method could be used in the future to determine how repeated uses of a removal tool on contaminated surfaces may affect removal efficacy. Similarly, future studies could investigate the possible spread of pathogens to the removal tools, and to additional surfaces with contaminated removal tools. There are also challenges with increasing scale of cement-based flooring, as households may experience deformations in their flooring, such as delamination and pop-outs (78, 79), which were not present on the lab-scale tile and create an uneven surface that may affect the survival of indicator organisms or pathogens.

Overall, the findings improve our understanding of *E. coli* survival on cement-based flooring and effective removal techniques. While environmental factors and soil presence are important to consider when evaluating the survival of *E. coli* and effective removal techniques, similarities in *E. coli* fate between the studied mix designs suggest the feasibility of using alternative mix designs, such as one that includes fly ash to offset the use of OPC, for flooring. In the future, the use of these alternative mix designs could be scaled up in large infrastructure projects, therefore reducing the embodied carbon in our built environments.

## Methods and Materials

### Cement-based tile preparation

Experiments were performed using cement-based tiles (127 mm squares, 12.7 mm depth) that mimicked household cement-based floors. Three mixes were used in this study; an Ordinary Portland Cement (OPC) concrete mix typically used in the United States, an OPC mortar mix typically used in Bangladesh, and an OPC mortar mix with 25% class F Fly Ash as a sustainable alternative (referred to as OPC fly ash mortar in this study). Both the OPC mortar mix and OPC fly ash mortar mix were used in the CRADLE trial. After mixing, tiles were cured in molds for 24 hours. They were then de-molded and wet-cured in a lime bath for 7 days, followed by air-curing for a minimum of 28 days. Mixing, casting, and curing followed ASTM C192 Standard Practice for Making and Curing Concrete Test Specimens in the Laboratory (80). Complete mix design information is available in the supplemental material.

All tiles used in experiments had a neat finish, as is typical in Bangladeshi household floors. To assess any variations in the finish roughness as a result of the different mix designs, roughness testing was performed with a mechanical stylus-type profilometer (Bruker 849 Dektak XT; Billerica, MA, USA). Three 2 cm long scan lines were averaged (using the arithmetic mean) to measure the roughness properties of each tile surface. The scan duration was 20 seconds and the stylus force was set to 3 mg for all the scans. The data were calibrated using the default setting in the profilometer software, which sets the ends of each measurement to zero. The profilometer measured the average roughness (the average deviation from the mean line), the root-mean-square roughness (more sensitive to peaks and valleys than the average deviation), and the total roughness height (the distance between the highest peak and lowest valley). Substantial variation in roughness between the tiles may affect *E. coli* survival and removal, as a rougher surface may shield the bacteria.

### *E. coli* preparation

*E. coli* K12 (ATCC 19853) cultures were prepared by inoculating 20 mL of tryptic soy broth (TSB; Becton, Dickinson, and Company, Sparks, MD, USA) with a loop of bacterial stock and incubating the culture at 37°C overnight. After incubation, the culture was assumed to be in the stationary growth phase (81) and was spiked onto the cement tiles without dilution. TSB was used to spike the *E. coli* onto the tile to replicate organic-rich matrices found in the field, such as mucus, saliva, vomitus, or feces (82).

### Experimental protocol

Multiple 2 cm wide squares were outlined on each cement-based tile and were sacrificially sampled. For the survival experiments, each 2 cm square represented a different time point while for the removal experiments, each square represented either a control or a removal activity sample. The three trials (biological replicates) were repeated on three separate tiles. Each square was spiked with either 50 μl of the *E. coli* culture and allowed to dry (which represents the no soil condition) or 50 μl of the *E. coli* culture mixed with 0.01 g of sandy loam (Ward’s Science, Rochester, NY, USA) and allowed to dry (representing the soil condition). Sandy loam is a type of soil made up of a blend of mostly sand with small amounts of silt and clay. Sandy loam is common throughout Bangladesh (83).

For survival experiments, following spiking, tiles were incubated at two environmental conditions - either 34°C and 75% relative humidity (RH) to mimic the wet season conditions (average of day and night) in Bangladesh or a 15°C and 75% RH to mimic the dry season conditions (average of day and night) in Bangladesh. The rainy season typically lasts from April to October, while the dry season ranges from November to February. The weather station closest to the CRADLE field site, located in Tangail, Bangladesh, had an average monthly maximum temperature of 33.6 °C in April and an average monthly minimum temperature of 10.1 °C in January (84). RH ranges from 69 - 85% throughout the year (85). RH conditions were maintained using a saturated salt solution (86). For tiles without soil at 34°C, samples were recovered and enumerated at 0, 0.33, 0.66, 1, 1.33, 1.66, and 2 hours. This specific timeframe was selected for this condition because in pilot experiments, *E. coli* without soil at 34°C decayed more rapidly than under other conditions and we aimed to capture the same number of time points before the *E. coli* reached our limit of detection (10 colony-forming units, CFUs). For all other experimental conditions (with or without soil at 15 °C and with soil at 34°C), the samples were recovered and enumerated at 0, 1, 2, 2.5, 3, 3.5, and 4 hours. Two to four hours represents a realistic length of time in the field in which the flooring would remain unperturbed since household members frequently enter or exit the home. Samples were recovered using a sterile cotton swab pre-wetted with sterile (autoclaved) phosphate-buffered saline (PBS; Fisherbrand, Waltham, MA, USA) and wiping across the square 10 times horizontally and 10 times vertically. Afterward, the swab was placed in a microcentrifuge tube filled with 1 mL of sterile PBS, vortexed for approximately 20 seconds, and then the *E. coli* in the PBS was enumerated using the plating technique described later in the Methods. Each experimental condition (3 mix designs x 2 environmental conditions x 2 with/without soil) was repeated for three trials.

For removal experiments, three common household activities were mimicked in the lab: mopping, sweeping, and walking. To simulate mopping, a sterile cotton swab (Fisherbrand, Waltham, MA, USA) pre-wetted with autoclaved deionized (DI) water was wiped across the square 5 times horizontally and 5 times vertically. To simulate sweeping, a sterile 9.5 cm long miniature straw broom (Healifty, Amazon Online) was swept across the square 5 times horizontally and 5 times vertically (see supplemental material, **Figure S1**). Walking experiments were simulated using 2 cm^2^ rubber tiles (X-Protector, Lisburn, Northern Ireland). Each rubber tile was pressed on the pre-marked square on the cement tile with a 500 g weight for 10 seconds to replicate a similar pressure from a one-year-old child walking around the household floor, which is approximately 130 g/cm^2^ (87) (see supplemental material, **Figure S1**). Each cotton swab, straw broom, or rubber tile was discarded after it was used to remove *E. coli* from the tile surface. *E. coli* were recovered from the tile using the same method as that used with the survival samples. Briefly, after the removal activities were completed, the remaining bacteria were recovered using a sterile cotton swab pre-wetted with sterile PBS which was wiped across the square 10 times horizontally and 10 times vertically. Afterward, the swab was placed in a microcentrifuge tube filled with 1 mL of sterile PBS and vortexed for approximately 20 seconds. *E. coli* in the resultant PBS sample was then enumerated using the plating technique described in the following Methods subsection. Each experimental condition (3 mix designs x 3 removal activities x 2 with/without soil) was repeated for three trials.

For each trial for both experiment types (i.e., survival and removal), a negative control was obtained by swabbing an area on the tile that was not spiked with *E. coli*. A positive recovery control was obtained for each trial of the removal experiments by swabbing a spiked square that did not receive any removal activity. The sample taken at time zero served as a recovery control for the survival experiments. Additional technical positive controls were run in the enumeration process for both experiment types by directly plating the *E. coli* culture without any cement-based tile interaction to ensure *E. coli* growth media was functioning.

### *E. coli* enumeration

Following sample collection, *E. coli* was enumerated using a spread-plating procedure. 50 μL of two or three dilutions of each sample (up to 1:10^7^ dilutions made using sterile PBS, based on preliminary experiments) were plated on tryptic soy agar (TSA; Becton, Dickinson, and Company, Sparks, MD, USA). Dilutions were plated in triplicate, samples were allowed to absorb for five minutes upright at room temperature, and the plates were incubated upside down at 37°C overnight. Colony Forming Units (CFUs) within the countable range (10 ≤ CFU ≤ 300) were counted and recorded the following day.

### Data analysis

Data analysis and calculations were performed in MATLAB (R2023a; The Math Works, Natick, MA, USA), Microsoft Excel (2018, Microsoft Corporation, Redmond, WA, USA), and R (2023, R Foundation for Statistical Computing, Vienna, Austria).

At each time point, CFUs within the countable range were averaged using the arithmetic mean. The 95% confidence intervals of the trials were calculated using the delta method. A first-order log-linear regression was used to model the behavior of surviving *E. coli* over time for each condition. The regression is described in Equation 1:

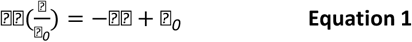

where *C*_*o*_ is the initial concentration of cells recovered from the tile, *C* is the concentration of surviving cells after an exposure time of *t* (minutes), *k* is the first-order decay rate constant (minutes^*−*1^), and *β*_0_ is the y-intercept. The models were evaluated by the *R*^2^ and p-values of the coefficients. A generalized linear model with a normal family and identity link was used on log-linear data (the y-dependent variable was transformed using logs, while the x-explanatory variable was unchanged). Regressions were performed for each experimental condition (12 regressions, 3 mix designs x 2 temperatures x 2 soil presences) to determine the average decay rate constant across each set of the three trials.

The removal efficiency was calculated using Equation 2. This equation calculates the percent removal of *E. coli* by dividing the average CFU concentration (*C*) after the removal activity by the initial average CFU concentration (*C*_*o*_) before the removal activity, subtracting that value by 1, and multiplying the fraction by 100%. The arithmetic mean of the percent removal was calculated using the three trials, and the 95% confidence intervals were determined using a t-test.

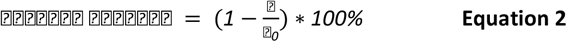

In the survival experiments, an *n*-way analysis of variance (ANOVA) was used to determine if mix design, environmental condition, or the presence of soil impacted the decay rate constant (*k*) of each condition.

The decay rate constants used in the ANOVA were a result of 36 linear regressions run for each condition and trial (3 mix designs x 2 temperatures x 2 soil presences x 3 trials). Regressions were run for the three trials within each experimental condition to provide sufficient degrees of freedom for the ANOVA. Interaction terms were not considered for the survival experiments. After the *n*-way ANOVA, a Tukey honestly significant post-hoc test was performed. Similarly, for the removal experiments in this study, the percent removal values of each condition were compared using an *n*-way ANOVA followed by a Tukey honestly significant post-hoc test to test any significance for the variables mix of design, removal activity, and presence of soil. Two-way interaction terms between removal activity and the presence of soil were considered. Shapiro-Wilks’ tests and Levenes’ tests verified that there was homogeneity of residuals and or variances and therefore the data could be reasonably approximated as normal, confirming the use of an ANOVA. All analyses were two-sided and were performed using a significance level of *α* = 0.05.

## Data Availability

Anderson, C.E., Hernandez, J., Hanif, S., Owens, L., Crider, Y., Billington, S.L., Lepech, M., Boehm, A.B., and Benjamin-Chung, J. (2024). Data set for: Evaluating the Survival and Removal of Escherichia coli from Surfaces Made with Traditional and Sustainable Cement-Based Materials in Field-Relevant Conditions. Version 1. Stanford Digital Repository. Available at https://purl.stanford.edu/jy612wv3964/version/. https://doi.org/10.25740/jy612wv3964.

## Acknowledgments

This work was supported by a grant from the Stanford Woods Institute for the Environment (281131) to J.B.C., the NSF Graduate Research Fellowships Program (GRFP) to C. A., and Air Force Institute of Technology Fellowship Program and John A Blume Earthquake Engineering Center Fellowship at Stanford University to J. H. J.B.C. is a Chan Zuckerberg Biohub Investigator. This study was performed on the ancestral and unceded lands of the Muwekma Ohlone people. We pay our respects to them and their Elders, past and present, and are grateful for the opportunity to live and work there.

